# The caddisfly gut microbiome contributes to the consumption of riparian leaf litter within rivers

**DOI:** 10.64898/2026.09.10.749941

**Authors:** Jonathan R. Dickey, Dahlia A. Loomis, Andrés Mauricio Caraballo-Rodríguez, Pieter C. Dorrestein, Sara L. Jackrel

## Abstract

Ecosystem functions can be regulated by biodiversity ranging from broad to narrow scales, including variation within a species. For most multicellular life, intraspecific variation is determined not only by an organism’s phenotype and genotype, but also variation imparted by their microbiome. Here, we investigate how leaf decomposition in rivers is affected by phenotypic variation in plant secondary metabolites (PSMs) and antibiotic-induced variation within the gut microbiomes of aquatic macroinvertebrate decomposers. We found that consumption by *Dicosmoecus* caddisflies was inhibited by the presence of dietary PSMs from riparian *Alnus rubra* trees, irrespective of the state of their gut microbiome. We further compared consumption of diets prepared from 20 different local and non-local *A. rubra* trees. We found that antibiotic-treated and control caddisflies consumed similar amounts of diets containing local *A. rubra* PSMs. However, compared to caddisflies with an intact gut microbiome, antibiotic-treated caddisflies consumed less of non-local *A. rubra* diets. Thus, decomposition by caddisflies could be regulated by both host adaptation to local resources and bacterial-mediated digestion of non-local resources. Overall, our study suggests that gut microbiomes can facilitate adjustment of macroinvertebrate decomposers to novel plant phenotypes, which may arise from shifting spatial distributions of plants in response to global change.

## BACKGROUND

The decomposition of leaf litter is a vital ecosystem function that provides nutrients to decomposer communities residing in terrestrial and aquatic environments.[1] Such ecosystems functions are often regulated by biological diversity at both broad and narrow taxonomic scales, including biological variation within a species.[2–4] Intraspecific variation is typically considered to be those traits caused by an individual’s phenotype and genotype, but variation imparted by host-associated microbes can be a third element of intraspecific variation with implications for host fitness, behavior, and population ecology.[5–8] The relative importance of these different aspects of intraspecific variation, as well as their interactive effects, on ecological interactions in natural systems still remains poorly understood.

In the context of plant-decomposer interactions, intraspecific variation in plant litter can affect the quality of litter resources for decomposers.[9–11] Litter variation includes differences in nutrient stoichiometry and the quantity of structural compounds, like lignin.[11] These aspects of litter quality may have fairly consistent effects for decomposers across environments. For example, high lignin content is detrimental to most decomposers, whereas high % N content is typically beneficial.[1,11,12] Litter quality can also vary in the types and quantities of plant secondary metabolites (PSMs), which are produced in the living plant as chemical defense strategies in response to pathogens and herbivores.[13] Beyond inhibiting the attackers of living plants, variation in PSMs has been shown to have cascading inhibitory effects on decomposers.[14] For example, intraspecific variation in PSMs of *Alnus rubra* was found to be one of the main regulating factors of leaf litter decomposition in both riparian soils and rivers.[15] Similarly, variation in PSMs among genotypes of *Arabidopsis thaliana* was found to have greater effects for decomposition rates than morphophysiological traits, such as leaf tensile strength and trichome density.[16] In comparison to the effects of structural components and nutrients in plant tissues, the contribution of PSMs to litter quality may be more context dependent on the composition and metabolic capabilities of decomposers inhabiting specific environments. A well-known example of this context dependency is the Home-Field Advantage phenomenon in which litter tends to decompose faster in the home environment than a geographically distant or ‘away’ environment.[17,18] This phenomenon has been documented most frequently at the species-scale, but it also occurs at the intraspecific scale within geographic regions where home and away plants may be separated by less than one kilometer.[19–21]

Plant-decomposer interactions may also be mediated by intraspecific variation within decomposers themselves. Invertebrate consumers have evolved adaptative means to counteract or even benefit from dietary PSMs.[22,23] Beyond genetic and phenotypic changes within these consumers, some invertebrates harbor a core microbiome within their gut that likely confers digestive functions, including the degradation of PSMs.[24–27] For example, two distantly related insects that feed on the same neurotoxic plant harbor similar core gut microbiota, suggesting that shared bacterial taxa may confer key detoxification functions.[28] Further, certain PSMs can inhibit herbivore digestion by reshaping the microbiota inhabiting their gut.[29,30] Thus, there may be important interactive effects between the composition of the invertebrate gut microbiome and intraspecific variation in PSMs that ultimately mediates feeding preferences among invertebrate consumers.

We investigate the independent and interactive effects of (1) intraspecific phenotypic variation in PSMs among riparian trees and (2) variation imparted by host-associated microbiota in a species of freshwater macroinvertebrate decomposer. We utilize the deciduous tree, *Alnus rubra*, which dominates many riparian zones of the Olympic Peninsula of Washington State, USA and provides sizable fluxes of greenfall into adjacent rivers during the summer growing season for freshwater macroinvertebrate decomposers.[19,31,32] Riverine decomposer communities in this region have also exhibited a persistent Home-Field Advantage, potentially as a result of geographic variation in *A. rubra* PSMs.[15] One of the largest and most abundant macroinvertebrates in these rivers is the larval caddisfly, *Dicosmoecus spp.* These dietary generalists are commonly observed grazing periphyton and consuming *A. rubra* leaf litter in rivers throughout this region.[33] Freshwater macroinvertebrates, such as *Dicosmoecus spp.*, are thought to be responsible for the largest proportion of leaf litter mass loss in riverine systems, contributing more to organic matter decomposition than bacterial and fungal decomposer communities.[34] Furthermore, prior work in this system has shown that *Dicosmoecus* larvae are sensitive to intraspecific variation in *A. rubra* leaf litter. Specifically, *Dicosmoecus* larvae fed *A. rubra* litter from their local riparian zone gained more body mass compared to those fed *A. rubra* litter from a non-local riparian zone.[33] Additionally, the *Dicosmoecus* core gut microbiome, which is dominated by Enterobacteriaceae, Clostridiales, Lactobacillales and Rhizobiales, was differentially affected in composition by feeding on local versus non-local populations of *A. rubra* litter, suggesting a potential role of the caddisfly gut microbiome in adjusting to the digestion of geographically variable *A. rubra* PSMs.[33]

In the present study, we first illustrate variation in local and non-local *A. rubra* PSMs across two riparian habitats, further documenting the geographic phenotypic variation in *A. rubra* defense compounds. We then test the effects of antibiotics on the composition and diversity of the *Dicosmoecus* larval gut microbiome. We hypothesize that antibiotic exposure will reduce the diversity of the caddisfly gut microbiome, with negative consequences for feeding rates, especially for those diets containing PSMs. To further understand the potential implications of the gut microbiome for host health and on rates of litter decomposition, we then assayed caddisfly feeding preferences for diets imbued with PSMs sourced from local versus non-local riparian populations of *A. rubra*. Based on prior work indicating negative fitness effects for *Dicosmoecus* restricted to a diet of non-local *A. rubra* [33], we hypothesized that caddisflies would prefer diets containing local PSMs. Further, given the probable role of the invertebrate gut microbiome in detoxification of PSMs [30], we hypothesized that this preference would be exacerbated among caddisflies with gut microbiomes that had been disrupted by antibiotics.

## METHODS

### Study sites

This study occurred in riparian zones of two third-order rivers of the Olympic Peninsula of Washington State, USA. Environmental parameters of the Pysht River and the Little Hoko River are described in **Table S1**. The riparian zones of both rivers are dominated by the deciduous tree, Red Alder (*Alnus rubra*), which is consumed by the dietary generalist, *Dicosmoecus sp*. As co-occurrence of the two species known to occur in Washington State, *D. gilvipes* and *D. atripes*, is rare, our study likely included individuals from only one species. We have observed *Dicosmoecus* individuals both grazing on periphyton and feeding on decaying leaves, which are more characteristic of *D. gilvipes* versus *D. atripes*, respectively.[35,36] Our past surveys of *A. rubra* individuals of these riparian zones have found that these two populations are similar in leaf stoichiometry (%N, %P, C:N, C:P, δ^13^C and δ^15^N), average leaf thickness, and tree trunk diameter (all analysis of variance model *p*-values > 0.10; *n* = 20).

However, as reported in detail elsewhere, *A. rubra* populations may vary in their composition of leaf PSMs, particularly the types and quantities of diarylheptanoids, ellagitannins and flavonoids.[15]

### Characterizing A. rubra leaf metabolomes via tandem liquid chromatography mass spectrometry

To determine whether *A. rubra* leaf PSMs are significantly different between these populations, we collected leaves from 10 individual *A. rubra* growing along the riparian zone of the Pysht River (local) and from 10 individuals growing along the riparian zone of the Little Hoko River (non-local). We hand-picked three green leaves from each tree that had no visible signs of herbivore or pathogen damage. Leaves were oven-dried for 48 hours at 60°C and ground into a fine powder using a mortar and pestle. Using 5 mg of this archived leaf powder, we extracted leaf metabolomes using 1 mL of 70% liquid chromatography–mass spectrometry (LC-MS) grade methanol spiked with 1 µM sulfadimethoxine on a Qiagen TissueLyser II (Qiagen, Hilden, Germany) set to 25 Hz for 5 minutes at ambient temperature. For data acquisition, 5 µL of extract was injected into the Vanquish ultra high-performance liquid chromatography system, which was coupled to a Q-Exactive quadrupole orbitrap mass spectrometer (Thermo Fisher Scientific, Bermen, Germany). See supplemental text for further details on sample extraction and instrument parameters.

Raw spectra were converted using the MSconvert plugin of ProteoWizard and imported into MZMine v.4.8.5 for MS1 and MS2 feature extraction.[37,38] Feature based molecular networking (FBMN) was conducted via GNPS2 (see GNPS2 task ID: 0370389e761e426c938fb4fe558a8ccc for full FBMN parameters).[39] Molecular fingerprinting and CANOPUS annotation was completed with SIRIUS v.6.3.0.[40–42] One local and three non-local trees were excluded from tandem LC-MS/MS due to insufficient archived leaf material to perform extractions. A full description of feature detection, molecular networking and analyses is provided in the supplemental text.

### Assaying caddisfly feeding preferences

#### Preparation of diet plates

To imbue artificial insect diets with PSMs from each of the 20 individual *A. rubra* trees (n = 10 per site), we used the leaf powder generated as described above to make metabolome extracts. From each tree, we extracted PSMs from 100 mg of ground leaf powder by mixing with 10 mL of 70% methanol in glass vials stored in the dark for 14 days.[43] We then transferred the supernatant containing the extracted PSMs to a new vial to evaporate. We then resuspended each PSM residue in 10 mL of sterile water (**Fig. S1**). Extracts from trees growing along the Pysht River were considered local and those from trees growing along the Little Hoko River were considered non-local relative to the origin of the *Dicosmoecus* larvae used in this study. We prepared all-purpose lepidopteran diets (Frontier Agricultural Sciences #F9772, Flemington New Jersey USA) either with or without the addition of *A. rubra* PSMs following our previously established methods.[15] See supplemental text for exact methodology.

#### Antibiotic treatment

To determine whether the gut microbiome plays a role in the feeding preferences of *Dicosmoecus,* we dosed larvae with an antibiotic cocktail to disrupt the gut microbiome composition and taxonomic richness. We incubated 100 caddisfly larvae that had originated from the Pysht River in plastic tanks filled with river water sourced from the Pysht. We maintained representative levels of dissolved oxygen (∼9.5 mg/L) recorded at the Pysht River in Summer 2022 using aquarium air pumps. Larvae were randomly assigned to either our control or antibiotic-treatment groups (n = 50 larvae per group). For our control group, larvae were maintained in Pysht River water and were fed the base diet prepared without antibiotics or PSMs. Larvae assigned to the antibiotic treatment group were maintained in Pysht River water treated with the following antibiotics: 10 mg/L of tetracycline, 10 mg/L of ciprofloxacin, 40 mg/L of streptomycin, and 4 mg/L of metronidazole. Antibiotics in the water were replenished every third day. Additionally, these larvae were fed base diet that were similarly prepared without any PSMs, but imbued with the following antibiotics: 1000 μg/g of tetracycline, 1000 μg/g of ciprofloxacin, 1000 μg/g of streptomycin, and 1000 μg/g of metronidazole. Antibiotic concentrations used to spike the water column and imbue diets were modified from [44] and [45]. Both control and antibiotic-treated larvae were maintained in these outdoor mesocosms under ambient lighting and temperature conditions for seven days at the University of Washington Olympic Natural Resources Center (ONRC; See **Fig. S2** for images of treatment mesocosms). To determine the effects of the antibiotic treatment on bacterial gut microbiomes, we froze 20 larvae from each group in individual sterile Whirl-Pak bags (Nasco, Fort Atkinson, WI, USA) at -20°C in the field and then at -80°C at University of California San Diego. The remaining 30 larvae per treatment group were then transferred to new containers with fresh antibiotic-free river water for 24 hours in order to clear antibiotics from their system prior to transfer to the *in-situ* river mesocosms.

#### In-situ river mesocosms

Larvae were then transferred to *in-situ* mesocosms in the channel of the Pysht River to determine feeding preferences between control diets, diets imbued with local *A. rubra* PSMs, and diets imbued with non-local *A. rubra* PSMs. Each mesocosm contained two 96-well plates for a total of 140 microwells filled with diet. Each microplate contained each of the 21 different diets: the control diet (n = 10 wells), each of the 10 *A. rubra* PSM-imbued diets from the Pysht River (local diet; n = 3 wells per tree), and 10 *A. rubra* PSM-imbued diets from the Little Hoko River (non-local diet; n = 3 wells per tree). See **Fig. S1** for an example 96 well plate containing randomly assorted larval diets. Each diet microplate was covered with a 4.75 mm mesh nylon seine netting that was secured with mini binder clips. This netting provided a surface for caddisflies to cling to during feeding, while also providing material that could be pinned under river stones to hold the diet plate in place. Each mesocosm contained either 10 larvae from the control group or 10 larvae from the antibiotic-treatment group. Once about half of the diet had been consumed across all mesocosms, the plates from each mesocosm were removed and sealed in plastic bags for transport, where we measured the quantity of diet consumed. See **Fig. S2** for images of the *in-situ* mesocosms and the supplemental text for our statistical approach.

### Characterizing the larval caddisfly gut microbiome

#### Larval dissections

Prior to dissection, we quantified size traits that may correspond with larval age class and the composition of their gut microbiome. We found that our control and antibiotic-treatment groups were statistically similar for each of these size metrics **(Fig. S3)**. This indicates that any differences observed in the gut microbiome could not be attributed to the physical size characteristics of the caddisflies assigned to each treatment group. We then followed surface sterilization methods and dissection techniques as described by [44] and in the supplemental text.

#### 16S rRNA sequencing of the larval gut

We extracted DNA from filters containing *Dicosmoecus* gut microbiomes of individuals from our control and antibiotic treatments that were maintained at the ONRC using the Qiagen DNeasy PowerSoil Pro Kit with a modified cell lysis and elution step. Extracted DNA was then snap frozen using liquid nitrogen and stored at -80°C. Library preparation and amplicon sequencing of the V4 region of the 16S rRNA gene were conducted at Argonne National Laboratory (Lemont, IL, USA) on an Illumina MiSeq instrument set to a PE150 cycle. Raw reads were processed with QIIME2 v.2022.8, the DADA2 plugin and phyloseq.[46–48] Merged reads were assigned bacterial taxonomy via SILVA 138 16S rRNA database.[49,50] We then rarefied to a sequencing depth of 28,484 reads using the *rrarefy* function within vegan v.2.6-8.[51] Visit the supplemental text for an extensive report on amplicon sequencing analyses.

## RESULTS

### Antibiotics alter the composition and predicted function of the Dicosmoecus gut microbiome

We found that the gut microbiomes of control versus antibiotic-treated caddisflies differed significantly in bacterial community composition using phylogenetically-based and non-phylogenetically based distance metrics (**Fig. 1A**; Bray-Curtis via db-RDA: *F*_1,35_ = 58.64, *p* < 0.001; see **Fig. S4** for db-RDA on uniFrac distances). Broad differences in bacterial community composition between these two groups are further illustrated in **Fig. S5**. The gut microbiomes of antibiotic-treated caddisflies were lower in alpha diversity compared to those from control larvae (**Fig. 1B**; Richness: *F*_1,35_ = 4.18; *p* < 0.05). This difference in alpha diversity between treatment groups remained even when considering only the most abundant bacteria (i.e., *q* = 2 in our Hill number analysis; **Fig. S6**). We also found greater heterogeneity in the composition of the gut microbiome within our antibiotic-treated group compared to the control group (beta-disper: *F*_1_ = 45.44, *p* < 0.001).

**Figure 1.**
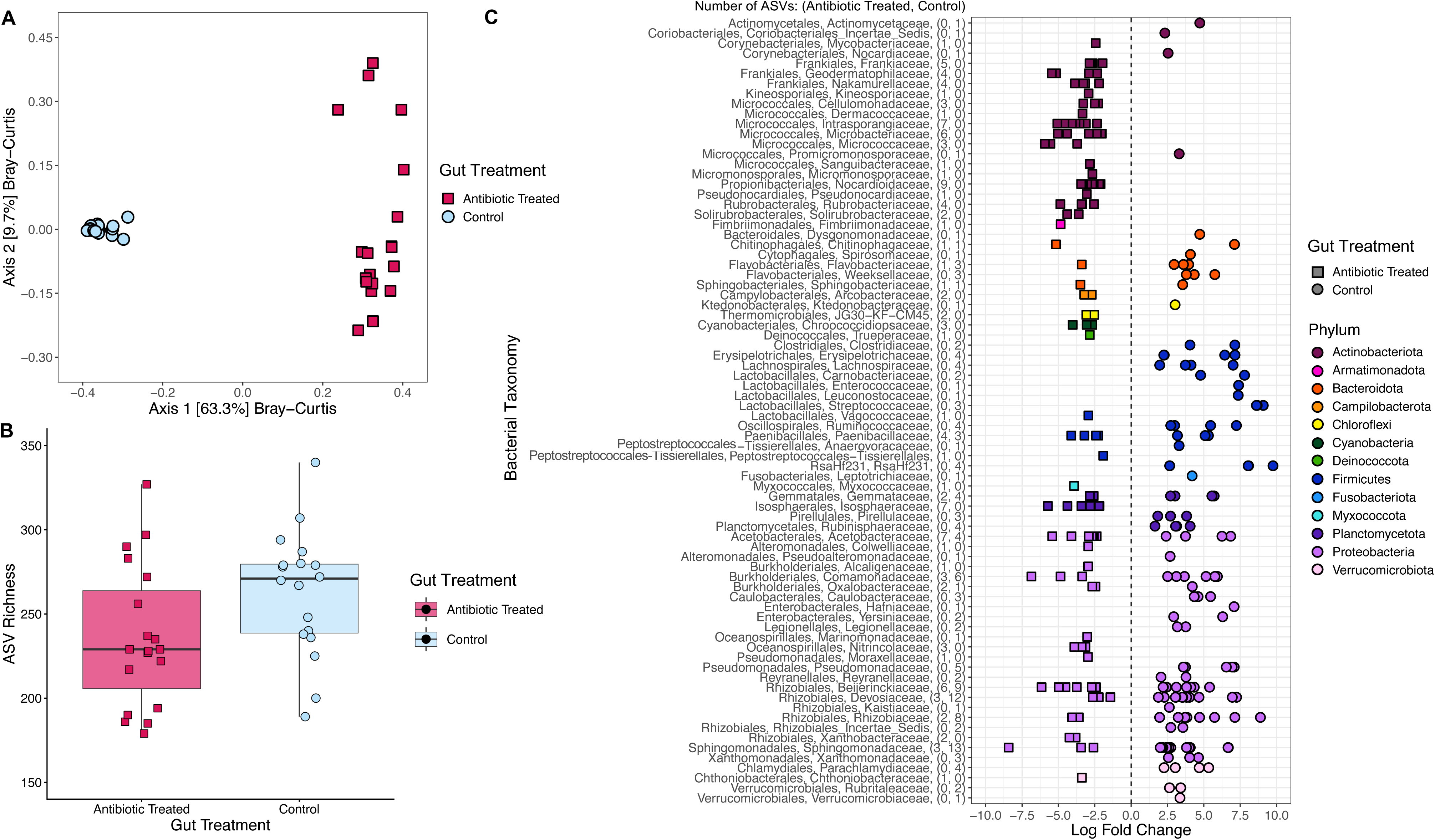
Community composition of the gut microbiome differed significantly between *Dicosmoecus* caddisfly larvae from the control versus antibiotic-treated groups. (**A**) Larvae treated with antibiotics experienced change in the composition of their gut microbiome relative to control larvae (db-RDA— *F*_1,35_ = 58.64, *p* < 0.001). (**B**) We observed that the gut microbiomes of control larvae were significant greater in taxonomic richness (ANOVA— *F*_1,35_ = 4.18; *p* < 0.05), indicating that antibiotic treatment had a negative effect on bacterial persistence within the gut. (**C**) Antibiotic treatment greatly reduced the Firmicutes in the larval gut microbiome, while Actinobacteriota increased in abundance relative to the gut microbiome of control larvae. Negative and positive log fold change indicate which taxa are over-represented in the gut microbiomes of antibiotic treated and control larvae, respectively. Parentheses on the y-axis represent the number of ASVs differentially abundant in antibiotic treated and control gut microbiomes. Differential abundance analysis was conducted with a significance level of p < 0.001 after false discovery rate correction.

In addition to these broad changes in community composition, we found that the gut microbiomes of our antibiotic-treated versus control caddisflies differed significantly in the relative abundance of 261 different ASVs belonging to families across 12 different phyla (**Fig. 1C**). For example, antibiotic-treated microbiomes were overrepresented in *Comamonadaceae* and *Sphingomonadaceae* of the *Proteobacteria,* while control caddisflies were overrepresented by *Firmicutes* (e.g., *Ruminococcaceae* and *Streptococcaceae*). Further, these compositional shifts in the caddisfly gut microbiome caused by the antibiotic treatment differentially affected predicted function among 165 functional pathways encompassing metabolic, cellular, environmental and genetic information processes (*p* < 0.05, **Fig. S7**). Several of these affected pathways could play roles in the degradation of *A. rubra* secondary metabolites, particularly those within the xenobiotic biodegradation and metabolism pathways (*p* < 0.05; **Fig. S8**).

Further, there were widespread predicted effects of antibiotic treatment on the capacity to metabolize amino acids, lipids and carbohydrates within the caddisfly gut (**Fig. S8**).

### Geographic variation in A. rubra leaf metabolites

The *A. rubra* leaf metabolome was comprised of 1,870 different PSMs and were dominated by certain PSMs in terms of high concentrations of individual features and total number of features. Specifically, high concentrations occurred among the phenylethanoids and diarylheptanoids (**Table S2**), whereas the ellagitannins were nested within the most numerically rich SIRIUS defined superclass, phenolic acids (n = 315). Furthermore, we found that local and non-local populations of *A. rubra* growing along the Pysht versus Little Hoko rivers strongly diverged in their leaf PSMs with divergence most driven by those features with Variable Importance in Projection (VIPs) scores > 1 (via PLS-DA on the total number of PSM features; **Fig. 2A**). Notably, 36.8% of total leaf PSMs contributed to population level discrimination between rivers. For example, the majority of discriminating features belong to SIRIUS defined superclasses such as the flavonoids, triterpenoids, and phenolic acids, where 29%, 27%, and 36% of features had site defining scores > 1 (**Fig. 2B**). Further, local and non-local metabolomes diverged in the relative abundances of discriminating compounds with VIPs > 1 (**Fig. 2C**).

**Figure 2.**
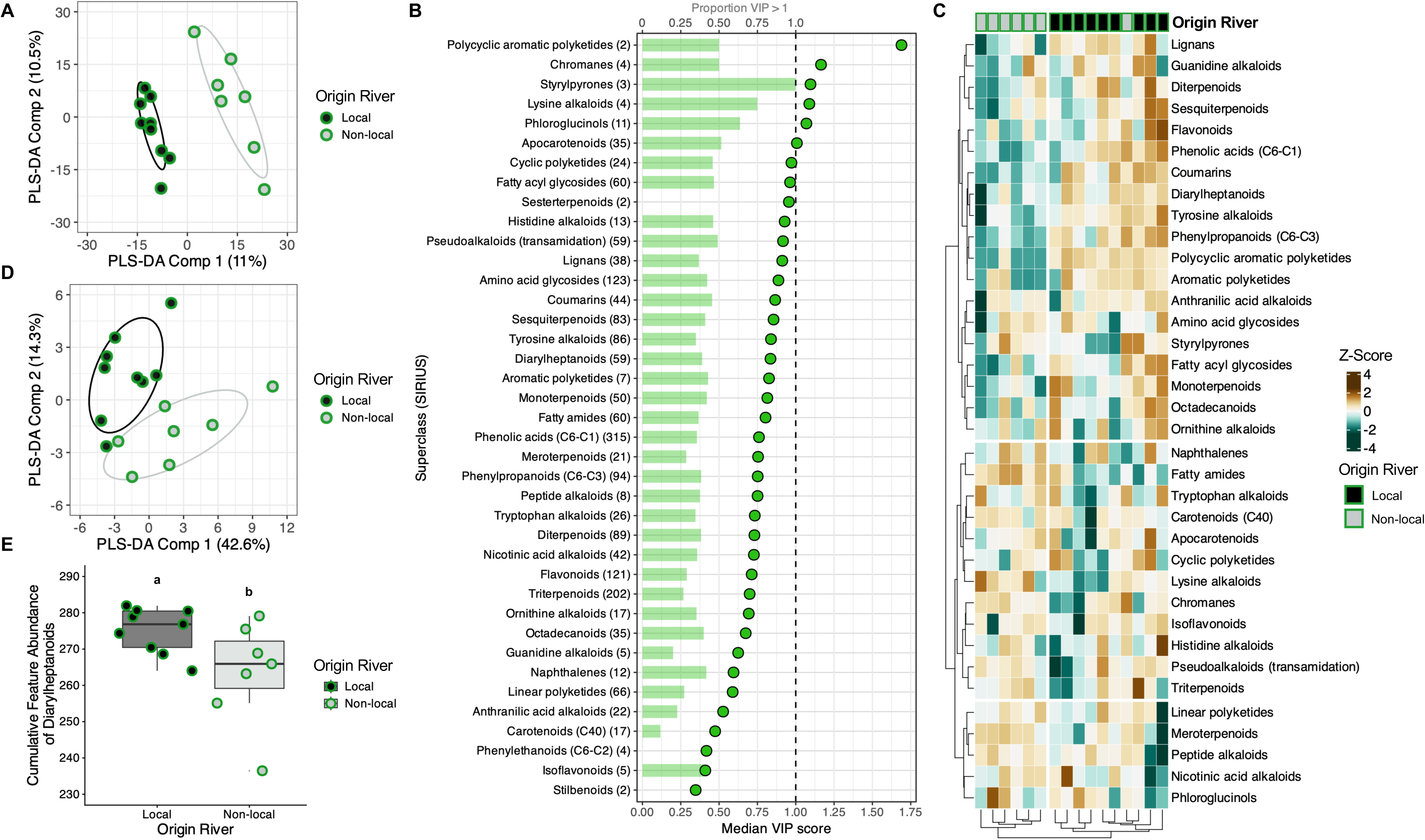
The leaf metabolomes of local and non-local populations of *Alnus rubra* diverged in their composition and abundance of plant secondary metabolites. (**A**) *A. rubra* from the Pysht (local) and Little Hoko River (non-local) diverge in their leaf metabolomes (PLS-DA; n = 1,870 PSMs). Ellipsoids represent one standard deviation of the multivariate score distribution. **(B)** Supervised clustering by origin river was driven by features with the largest Variable Importance in Projection (VIP) scores. SIRIUS defined PSM superclasses are sorted by the median VIP scores (lower x-axis) with the number of features per PSM superclass listed in parentheses. In addition, the proportion of features with VIP scores > 1 are represented by shaded bars and the upper x-axis. This illustrates which PSM superclasses contribute above average towards origin river discrimination. While diarylheptanoids and ellagitannins (nested within the phenolic acids C6 - C1 superclass) features do not have the largest VIP scores on average, 38.98% and 35.56% of their features have VIP scores > 1, respectively. **(C)** PSMs with VIP > 1 show differences in relative abundance between *A. rubra* populations. Specifically, in local *A. rubra* populations we observed a higher representation of lignans, flavonoids, diarylheptanoids and phenylpropanoids among other notable PSM superclasses. **(D)** As diarylheptanoids have previously been documented to vary among *A. rubra* populations, we conducted a PLS-DA with features assigned to this superclass and similarly show distinct population level clustering. In addition, **(E)** the cumulative abundance of Diarylheptanoids in fresh leaves (VIP > 1) is significantly different among *A. rubra* populations (ANOVA— Origin River: *F*_1,14_ = 4.90, *p* = 0.04).

Specifically, the leaf metabolomes of local *A. rubra* tended to contain greater abundances of lignans, flavonoids and diarylheptanoids. We then preformed PLS-DA on PSMs superclasses that we expected to vary geographically. Specifically, we found that local and non-local *A. rubra* populations differed when considering only the composition and relative abundances of diarylheptanoids (**Fig. 2D-E**; diarylheptanoids with VIPs > 1; ANOVA— Origin River: *F*_1,14_ = 4.90, *p* = 0.04). In contrast, these populations did not significantly differ among either ellagitannins or flavonoids, although PLS-DAs still revealed distinct compositional clustering (ANOVAs— ellagitannins: *F*_1,14_ = 0.21, *p* = 0.65; flavonoids: *F*_1,14_ = 0.06, *p* = 0.81; **Fig. S9**).

See **Table S3** for the top ten most discriminating compounds between the local and non-local *A. rubra* populations for each PLS-DA analysis.

### Interactive effects of A. rubra PSMs and the gut microbiome on caddisfly feeding preferences

Caddisfly larvae consumed on average 27.3% less of the diets imbued with *A. rubra* PSMs compared to the control diet prepared with just water (**Fig. 3**; linear mixed effects model (LMER)— *F*_5,27.4_ = 4.11, *p* = 0.007). This preference for the control diet was observed regardless of whether the caddisfly gut microbiome had been disrupted with antibiotics (Planned contrasts via LMER— control larvae: *t*_12.3_ = 3.36, *p* = 0.003, antibiotic treated larvae: *t*_12.3_ = 3.28, *p* = 0.006). Among the 20 diets imbued with *A. rubra* PSMs, caddisflies fed on the different diets at varying rates, ranging from 102.2 mg ± 7.78 SE consumed for the most preferred diet to 70.8 mg ± 6.64 SE consumed for the least preferred diet. Overall, there was no systematic preference for those diets imbued with PSMs from the local Pysht *A. rubra* population compared to those diets imbued with PSMs from the Little Hoko *A. rubra* population (Planned contrasts of local vs non-local *A. rubra* diets via LMER: *t*_18_ = -0.57, *p* = 0.58; local diets: *µ* = 81.83 mg ± 2.08 SE; non-local diets: 85.28 mg ± 2.20 SE).

**Figure 3.**
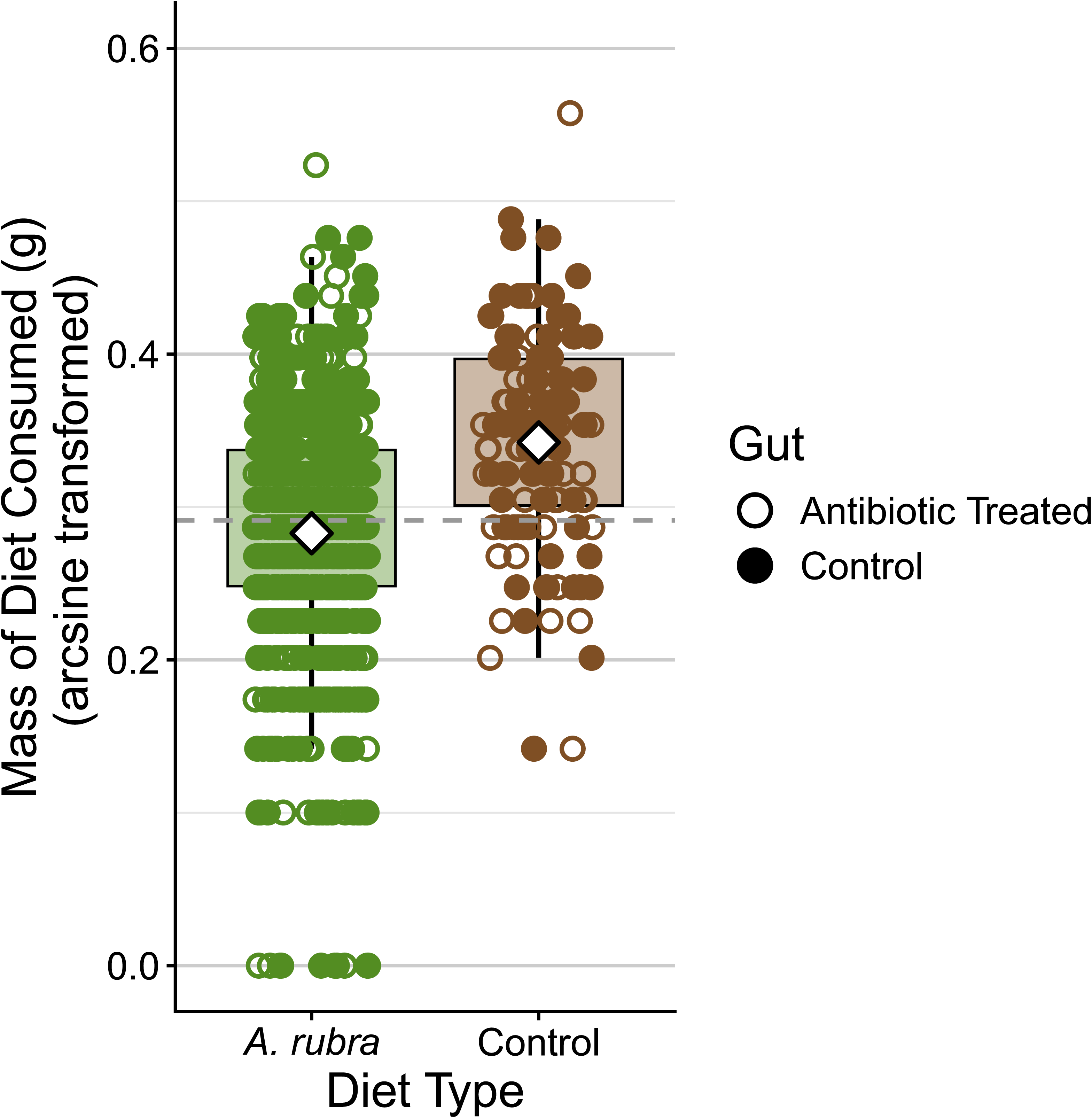
Diets imbued with *A. rubra* leaf metabolites were consumed less frequently by caddisflies compared to control diets. *Dicosmoecus* caddisfly larvae preferred to consume simple all-purpose lepidopteran diet over diets imbued with complex aromatic PSMs extracted from fresh *A. rubra* leaves (LMER— *F*_5,27.4_ = 4.11, *p* = 0.007). This preference for control diet was observed regardless of whether the caddisfly gut microbiome had been disrupted with antibiotics or not (Planned contrasts via LMER— control larvae: *t*_12.3_ = 3.36, *p* = 0.003, antibiotic treated larvae: *t*_12.3_ = 3.28, *p* = 0.006). Dashed gray line represents the global mean diet consumed (µ = 0.291 g). White diamonds represent the mean mass of diet consumed for each diet type.

However, the condition of the larval gut microbiome affected consumption rates of diets prepared with different sources of *A. rubra* PSMs. Specifically, while antibiotic-treated and control caddisfly larvae consumed similar quantities of diets imbued with local *A. rubra* PSMs, antibiotic-treated larvae consumed on average 7.14% less of diets imbued with non-local *A. rubra* PSMs (**Fig. 4A-B**; planned contrasts via LMER— antibiotic-treated vs. control larval consumption of local diets: *t*_814_ = -0.01, *p* = 0.99; non-local diets: *t*_814_ = 1.98, *p* = 0.047). This reduced consumption by the antibiotic-treated larvae was observed for nine of the ten different non-local *A. rubra* diets (**Fig. 4C**). The disruption of the gut microbiome by antibiotics had no effect on the more palatable control diet throughout their daily feeding (planned contrasts via LMER—control diets: *t*_814_= 1.02, *p* = 0.31). See **Fig. S10** and **Fig. S11** for consumption patterns of non-local and local diets, respectively. Despite widespread suppressed consumption of the non-local diets by antibiotic-treated caddisflies, control and antibiotic-treated caddisflies tended to exhibit similar mean ranked preferences for the various *A. rubra* diets (**Fig. 4D**; control diet plotted for comparison but not included in statistics: Spearman’s rank-correlation: ρ = 0.45, *p* = 0.045).

**Figure 4.**
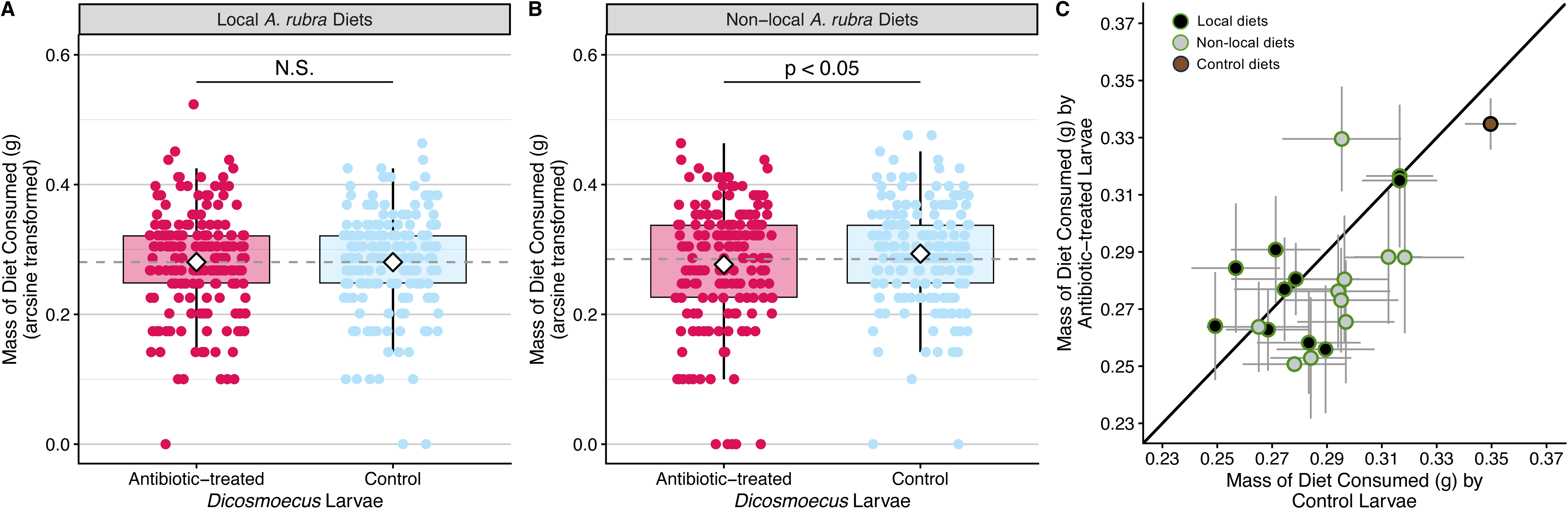
Diets imbued with non-local *A. rubra* leaf metabolites were consumed less frequently by antibiotic treated larvae than control caddisflies. Antibiotic-treated and control *Dicosmoecus* caddisfly larvae consumed similar quantities of **(A)** diets imbued with PSMs of local *A. rubra* trees from the Pysht River population (Planned contrasts via LMER— *t*_814_ = -0.01, *p* = 0.99). In contrast, **(B)** diets that were imbued with non-local *A. rubra* PSMs were consumed significantly less by antibiotic-treated larvae with a disrupted gut microbiome (Planned contrasts via LMER— *t*_814_ = 1.98, *p* = 0.047). Dashed gray lines depict mean consumption for each diet type, irrespective of caddisfly group (Local: µ = 0.280 g; Non-local: µ = 0.285 g). White diamonds depict mean consumption by each caddisfly group. **(C)** Of the ten different diets prepared with the PSMs from non-local *A. rubra* trees from the Little Hoko River population, nine tended to be consumed less by antibiotic-treated larvae than the control larvae, as represented by grey-filled points falling beneath the 1:1 line (mean mass consumed ± 1 SE, see Fig. S10 for illustration of raw data). In contrast, of the ten different diets prepared with PSMs from local *A. rubra* trees from the Pysht River population, only four were consumed less by antibiotic-treated larvae, as depicted by black-filled points falling beneath the 1:1 line (see **Fig. S11** for illustration of raw data).

## DISCUSSION

Intraspecific variation in the PSM signature of leaf litter can alter the fitness of decomposers.[33,52,53] Decomposer communities have often responded to this selective pressure in their food resources by adjusting or adapting to local PSMs, resulting in a Home-Field Advantage. However, for caddisflies and other macroinvertebrate decomposers, it is unknown whether this adjustment to intraspecific variation in PSMs arises from adaptations within the host organism or from the microbiota inhabiting the host gut. We demonstrated that the presence of dietary *A. rubra* PSMs deterred feeding by caddisflies in comparison to diets lacking these PSMs. This inhibitory effect of *A. rubra* PSMs was similar for those caddisflies with and without intact gut microbiomes, suggesting that the gut microbiome does not appear broadly essential for the consumption of all *A. rubra* PSMs. Furthermore, we found that intraspecific variation among *A. rubra* PSMs had substantial consequences for caddisfly feeding preferences. When honing in on the geographic origins of this intraspecific variation in *A. rubra* PSMs, we found that an intact gut microbiome appeared to facilitate the consumption of more novel diets comprised of non-local *A. rubra* PSMs. Specifically, we observed reduced consumption by the antibiotic-treated larvae relative to the control larvae for 90% of the non-local *A. rubra* diets. Overall, these results suggest that the gut microbiome might not be critical for the consumption of a local diet that a consumer has been able to adapt to over time. In contrast, the gut microbiome may play a more important role in facilitating host adjustment to novel food resources. Such novel food resources may arise more frequently in the Anthropocene as plant populations shift across the globe in response to human activities, potentially resulting in plant-decomposer mismatches.

### Antibiotics disrupt the composition and probable function of the caddisfly gut microbiome

As invertebrates are incredibly diverse and occupy a wide array of ecological niches, their gut microbiomes can vary substantially in taxonomic composition, diversity and function.

We found that the gut microbiome of *Dicosmoecus sp.* was similar in composition to that of many insect taxa, with a predominance of *Proteobacteria* and *Firmicutes*, and specifically to that of other detritivores in the prevalence of *Clostridiales* and *Bacteroidales*.[54–56] Further, we found that antibiotics had varied effects on these microbial inhabitants comprising the caddisfly gut microbiome. This result was expected as insects have been studied as reservoirs for antimicrobial discovery due to frequent exposure of their gut inhabits to an array of dietary PSMs, many of which have antimicrobial properties.[57] Specifically for caddisflies, many of the ellagitannins and diarylheptanoids that comprise *A. rubra* litter are known to have antibacterial properties.[58–60] Thus, these variable responses of bacterial taxa that comprise the gut microbiome of caddisflies could be due in part to evolved resistance to antimicrobial compounds that occur naturally in the caddisfly diet.

Beyond an overall decrease in richness within the caddisfly gut, we found that antibiotic treatment had variable effects for and against certain bacterial phyla, with likely cascading implications for metabolic function. The *Actinobacteriota* were overrepresented in larvae treated within antibiotics, perhaps due to the prevalence of this phylum in producing antibiotics and antibiotic-resistance genes as a self-protecting mechanism against these compounds.[61] This increased abundance of the *Actinobacteriota* of antibiotic-treated caddisflies likely explains the overrepresentation of predicted functional pathways related to the biosynthesis of secondary metabolites, including phenylpropanoids, isoflavonoids, and terpenoids.[62] While production of these secondary metabolites is largely unique to plants, a few exceptions have been found in the *Actinobacteriota.*[63] In contrast, the *Firmicutes* had the opposite response and likely the lowest tolerance to antibiotic exposure, perhaps specifically due to tetracycline, which was shown to eliminate *Firmicutes* in adult fly gut microbiomes.[64,65] The most abundant phyla inhabiting caddisfly gut microbiomes, the *Proteobacteria*, had a more variable response to antibiotics than other phyla, where some families increased in relative abundance, while others were unaffected or experienced only moderate declines upon exposure. Reductions in *Firmicutes* and certain *Proteobacteria* caused by antibiotic treatment could be implicated in reduced capacity of caddisflies to consume non-local *A. rubra* PSMs, as these phyla contain several taxa that are capable of aromatic PSM catabolism in anaerobic environments such as the insect gut (i.e., the *Pseudomonadaceae*, *Streptococcaceae*, *Lachnospiraceae*).[66] Further, the intact gut microbiome was enriched in predicted functions relating to xenobiotic degradation, particularly aromatic compounds with structural similarity to PSMs (e.g., nitrotoluene, styrene, xylene), which suggests that the intact *Dicosmoecus* gut microbiome may generally be adept at cleaving the aromatic ring structures that comprise most PSMs.[67]

### The caddisfly gut microbiome facilitates consumption of non-local A. rubra PSMs

Caddisflies appear to rely on their gut microbiome when challenged with novel dietary PSMs. Specifically, we found that caddisfly larvae treated with antibiotics tended to consume less of diets prepared with non-local *A. rubra* PSMs. In contrast, those with an intact gut microbiome showed no systematic preference between local versus non-local diets. This lack of preference among caddisflies with an intact gut microbiome was not unexpected and still mirrors the Home-Field Advantage phenomenon since some *A. rubra* within a single population have traits associated with high toxicity and slow rates of decomposition whereas others have more palatable traits.[15] Thus, while local plants do not always decompose at a faster rate than non-local plants, the Home-Field Advantage occurs when local plants breakdown relatively faster *than expected* based on their palatability traits, which can be demonstrated through reciprocal transplant studies.[18] The difference that we observed in dietary preferences between caddisflies with and without an intact gut microbiome suggests that the caddisfly host may have some adaptive genetic mechanism for tolerating PSMs found in abundance in their local diet, but may then rely on their gut microbiome when challenged with novel dietary PSMs.

The increased role of bacterial metabolism in the breakdown of non-local PSMs within the caddisfly gut parallels prior results focused on the bacterial communities that inhabit and decompose leaf packs of *A. rubra* in these river ecosystems. Specifically, leaf packs of non-local *A. rubra* retain a specialized community of *Burkholderiales* bacteria, which are known for their capacity to cleave aromatic rings in aerobic environments, for a longer period of time than leaf packs of local *A. rubra*, which instead rapidly shift to a generalist community of bacterial decomposers.[68] Thus, while this longer residence time of *Burkholderiales* may facilitate eventual breakdown of aromatic rings, these non-local leaves ultimately decompose relatively more slowly than expected, resulting in a Home-Field Advantage. Similarly, even with bacterial-mediated degradation of dietary PSMs in the intact caddisfly gut microbiome, long-term restriction to a diet of non-local *A. rubra* was shown to slow growth rates among *Dicosmoecus*.[33] This suggests that non-local PSMs may require more intensive breakdown by bacteria, both within the gut of caddisflies and free-living in the decomposing leaf matrix.

Reduced consumption of non-local *A. rubra* PSMs requires that *Dicosmoecus* are capable of distinguishing among diets and behaviorally shifting their feeding preferences. Such behavioral shifts induced by antibiotic exposure have been documented in other insects in which the gut microbiome has been implicated in behavioral regulation of the host. Specifically, diet-induced shifts in the gut microbiomes of the cockroach *Blattella germanica* have been shown to alter the host’s olfactory responses towards different food resources.[69] Further, our functional predictions for the antibiotic-induced shift in the gut microbiome of *Dicosmoecus* suggests that significant changes in the production of indole could be an underlying mechanism for this behavioral shift. Gut bacteria often produce large quantities of indole within the animal digestive tract where it acts as an important interkingdom signaling molecule through interaction with intestinal epithelial cells and ultimately modulation of behavior and olfaction.[70,71] For example, *Lactobacillus* within the honeybee gut microbiome was shown to promote olfactory learning and memory behavior via the production of indole derivatives.[72] Future directions could therefore explore how specific taxa comprising the caddisfly microbiome might not only modulate the breakdown of aromatic PSMs, but also the behavioral aspects that control *Dicosmoecus* feeding preferences.

### Limitations, future directions and conclusions

One important limitation of this study was the necessity for destructive sampling of the gut microbiome, which required that we analyze effects of the antibiotic treatment on the microbiome composition of one subset of caddisflies, whereas the remainder were used in feeding assays. Further, scaling up diet studies such as these to include multiple populations of local and non-local *A. rubra*, as well as a full reciprocal transplant design and collecting individuals at the conclusion of feeding trials, may further elucidate interactive effects between the gut microbiome, PSM origin and larval feeding behaviors. Future work should also evaluate the resilience of the gut microbiome in compositional structure and function following a disturbance of antibiotic exposure and subsequent return to the natural environment.

Additionally, predictions for bacterial function were inferred from 16S rRNA marker gene surveys with the PICRUSt2 tool, which makes predictions of gene content in an environmental sample by referencing metagenomic databases rather than actual sequencing of metagenomes.[73] Accurate predictions require that closely related reference genomes exist for each taxon found in an environmental sample, which is less likely for non-model organisms in understudied ecosystems.[73] Future studies could more directly uncover functional consequences of antibiotic treatment on the caddisfly gut microbiome by employing metagenomic and metatranscriptomic approaches. Lastly, to more fully evaluate how plant- decomposer interactions may shift in the midst of global changes, future studies should track whether antibiotic treatment, paired with exposure to a greater number of diets spanning more spatially variable litter chemistry, has long-term fitness consequences for invertebrate decomposers.

Overall, identifying how gut microbiomes regulate the feeding behavior of freshwater macroinvertebrate decomposers could shed light on the role of host-associated bacteria in regulating nutrient and energy transfer through riverine food webs. Our work elucidates how multiple elements of diversity at the intraspecific scale can regulate the important process of leaf litter decomposition. Given declines and shifting distributions of biological diversity during the Anthropocene, this work clarifies how changes in diversity patterns of plants and bacteria, even at relatively small taxonomic and spatial scales, has the potential to have larger implications for important ecosystem functions.

## ACKNOWLEDGEMENTS

We thank Dylan Meyer for his assistance in completing the caddisfly feeding assays and collecting field data. We thank J. Murray, Merrill & Ring Inc., and Hoko Rivers State Park for permitting research on their lands.

## DATA AVAILABILITY

All fastq files associated with the 16S rRNA dataset have been made publicly available in NCBI Short Read Archive under the BioProject ID PRJNA1222423. RScripts, MZmine batch file, and the SIRIUS annotation settings can be found at the following GitHub repository https://github.com/jrdickey9/Caddie_Antimicro_Diets. Full information for the LC-MS/MS protocol is available in the method file, provided as part of the public dataset MSV000099843 in the MassIVE repository (https://massive.ucsd.edu/).

## DECLARATION OF AI USE

We did not use AI-assisted technologies in creating this article.

## FUNDING

This project was supported by funding from UC-San Diego startup funds and the National Institutes of Health NIGMS R35GM142938 to S.L.J. A.M.C-R. and P.C.D. were supported by the Gordon and Betty Moore Foundation, GMBF12120 and https://doi.org/10.37807/GBMF12120

## CONFLICTS OF INTEREST

P.C.D. is an advisor and holds equity in Cybele, BileOmix and Sirenas and a Scientific co- founder, advisor, holds equity and/or received income to Ometa, Enveda, and Arome with prior approval by UC-San Diego

## AUTHOR CONTRIBUTIONS

J.R.D: data curation, formal analyses, investigation, methodology, project administration, supervision, validation, visualization, writing—review and editing. D.A.L.: methodology, formal analyses, visualization, writing—original draft. A.M.C-R.: funding acquisition, methodology, validation, writing—review and editing. P.C.D.: funding acquisition, resources, writing—review and editing. S.L.J.: conceptualization, funding acquisition, resources, supervision, validation, writing—review and editing.

