## Supplemental Information File for "The caddisfly gut microbiome contributes to the consumption of riparian leaf litter within rivers"

**TITLE**

**SUPPLEMENTAL TEXT**

***A. rubra metabolome extraction, acquisition and LC-MS/MS instrument parameters***

Fresh leaves were collected from each *A. rubra* tree and were dried for 72 hours at 60°C and then ground into a fine powder using a mortar and pestle. We extracted leaf secondary compounds with 5 mg of dried tissue and 1 mL of 70% LC-MS grade methanol imbued with 1 µM of sulfadimethoxine (extraction internal standard) using a Qiagen TissueLyser II (Qiagen, Hilden, Germany). TissueLyser II parameters were set to 25 Hz for 5 minutes at room temperature. To remove leaf material, we centrifuged extractions at 15,000 rpm for 5 minutes at room temperature and then transferred 400 µL of the supernatant to a 96 well deep plate. Sample concentration was then conducted with a Labconco centrivap system for 24 hours at 25°C. Prior to sample injection, samples were resuspended with 200 µL 70% LC-MS grade methanol with 1 µM sulfachloropyridazine as an internal standard.

Chromatographic separation occurred with a C18 core-shell column (Kinetex, 150 x 2.1 mm, 1.7 µm particle size, 100 Å pore size, Phenomenex, Torrance, USA) with a flow rate of 0.5 mL/min. Solvent A was water with 0.1% formic acid (FA) and Solvent B was acetonitrile with 0.1% formic acid. After injection, extractions were eluted via a linear gradient with the following specs: 0 – 8.5 min at 5% Solvent B, 8.5 – 10.5 min at 5-99% Solvent B, 10.5 – 11.5 min at 99% Solvent B, 11.5 – 12.75 min at 5% Solvent B, 12.75 – 14 min at 99% Solvent B, followed by a 1 min washout phase at 99% B. MS/MS spectra were obtained in positive ion data dependent acquisition (DDA) mode. Electrospray ionization parameters were 52 L/min sheath gas flow, 14 L/min auxiliary gas flow, 0 L/min sweep gas flow with auxiliary gas temperature set to 400°C. Spray voltage was 3.5 kV with the inlet capillary temperature set at 320°C. Full MS scan range was 100 – 1100 m/z at a resolution of m/z 200 (R_m/z 200_) of 35,000 with one micro-scan. Maximum injection ion time (IT) was 100 ms with an automated gain control (AGC) target of 5.0E5 for survey scans and MS/MS scans. In DDA mode, up to 5 MS/MS spectra per micro-scan were recorded with R_m/z 200_ = 17,500 with one micro-scan. Normalized collision energy increased from 20, 30, to 40% with the default charge state set to 1. MS/MS scans were triggered within 2 – 15 seconds of their first occurrence at their apex chromatographic peaks. Dynamic exclusion was set to 10 seconds. Ions with unassigned charge states and isotope peaks were excluded from DDA. Lastly, technical blanks, quality controls (QC mix of sulfamethazine, sulfamethizole, sulfachloropyridazine, sulfadimethoxine, amitriptyline, and coumarin-314 each at 10 µM), and a QC sample pool were injected every 10 samples to monitor LC-MS/MS performance.

***Feature detection and molecular networking***

Raw spectra were converted with the MSconvert plugin from ProteoWizard and then MS1 and MS2 features were extracted with MZMine v.4.8.5.[1,2] MS1 and MS2 detection was completed using the factor of the lowest signal (noise factors = 4.0 and 2.0, respectively). We set chromatogram building parameters to 4 minimum consecutive scans, 1.0E4 minimum absolute height, and 20 ppm m/z tolerance. We then split chromatograms using the local minimum feature resolver with these parameters: 80% chromatographic threshold, 0.075 minimum search range retention time and 1.8 minimum ratio of peak top/edge with a minimum of 4 scans. Next, we applied the 13C isotope filter and finder with the following join aligner parameters: 85% weight for m/z and a 0.35 retention time tolerance. Gap filling was conducted and then we filtered for duplicate peaks. Next, ion identity networking was completed with the metaCorrelate feature grouping algorithm. The final feature table and spectral lists were exported from MZMine with the GNPS and SIRIUS export functions. GNPS2 feature-based molecular networking (FBMN) with library analog search was conducted with 0.05 for precursor and fragment ion tolerance.[3] Network parameters included minimum cosine score of 0.7, minimum 5 matching peaks and networking max shift set to 1999, Top K = 10, and 100 max component size. FBMN library search parameters were minimum cosine set to 0.7, minimum 5 matching peaks. Chemical compound annotations from GNPS2 libraries were considered level 2 (cosine score > or equal to 0.7 and > or equal to 5 matched peaks) while predictions from SIRIUS were considered level 3 according to the Metabolomics Standard Initiative.[4,5]

We then exported the MZMine feature table along with annotations from FBMN and SIRIUS for further processing, normalization and analyses in R. Total ion current (TIC) was evaluated to determine successful sample injection and presence of sulfadimethoxine. Upon successful identification, the feature corresponding to sulfadimethoxine was filtered out of feature and annotation tables. Features found within QC mix and blanks with an average intensity of 5 times or greater within QC sample pools were removed. We then employed HomologDiscoverer to identify contaminant features found in any homolog runs or correlation groups with the following SIRIUS annotations: polyethylene glycols (PEGs) and common lab solvents, surfactants and contaminants.[6] We then identified and removed features with a positive or negative deltamz indicative of a change in mass of a PEG polymer from a previous recorded contaminant (≥ 44.02 and ≤ 44.03 or ≥ 88.05 and ≤ 88.06).

***Feature normalization, intensity filtering and statistical analyses***

Blanks, QC mix, and QC sample pools were removed from feature and annotation tables. Features intensities were then normalized using the internal standard, sulfachlorpyridazine. We first calculated the coefficient of variation for sulfachlorpyridazine across all samples and identified sample outliers that were excluded from the global mean calculation. We then normalized feature intensities by dividing all feature intensities by the per-sample internal standard intensity and multiplied this value by the internal standard’s global mean intensity. Features detected in less than four samples were removed. To remove low intensity noise, we filtered feature tables to retain only those with a maximum intensity > 20,000. Features with no SIRIUS annotation at the superclass level were removed. Prior to analyses, we stabilized variance of feature intensities by performing a log transformation.

To discern variation among PSMs from trees originating from the Little Hoko and Pysht Rivers, we performed partial least squares discriminant analysis (PLS-DA) on 1) the total number of leaf PSMs and 2) key compound groups (e.g., diarylheptanoids, ellagitannins and flavonoids) with the plsda function of the mixOmics package v.6.30.0.[7] Diarylheptanoid features were characterized via SIRIUS superclass annotation and verified by manual examining parent molecular mass and fragmentation patterns for each candidate feature as previously described.[8,9] Given that ellagitannins were nested within the phenolic acid (C6 – C1) SIRIUS superclass, we searched both SIRIUS and FBMN compound annotations matching “ellag”, “ellagic”, or “ellagitannin” and then similarly verified these features given parent molecular mass and fragmentation patterns. Candidate flavonoid features were subset based on successful SIRIUS superclass annotation. Features with zero or near-zero variance across samples were removed with *nearZeroVar* function within the caret v.7.1.0 package.[10] PLS-DA outliers were determined by comparing per sample squared Mahalanobis distance to the 90% quantile of the Chi-Squared distribution. We then calculated the variable importance in the projection (VIP) score for each feature, where scores greater than 1.0 were considered as features driving group separation in each PLS-DA. To test for geographic differences in the relative intensities of discriminating PSMs, feature intensities for VIPs were summed per superclass PLS-DA for each *A. rubra* tree. Linear models were built where cumulative feature abundances were predicted by river (local vs non-local) as a main effect. To illustrate geographic variation in total leaf PSM composition, we subset our feature table for VIP > 1 and built a heat map using the *comp_heatmap* function in microViz v. 0.12.6 package (seriation completed with Ward’s Optimal Leaf Ordering based on Euclidean distances).[11]

***Larval diet preparation at the Olympic National Resources Center***

The base diet was comprised of 10.5% of the supplied agar, 10.5% agarose (Sigma Aldrich), and 79% of the dry Frontier Agriculture Sciences diet. By using both agar and agarose as gelling agents, we minimized the disintegration of diets upon submergence in river water. For the control diet with no *A. rubra* PSMs, we mixed 10 g of the warmed base diet with 3.3 mL of water. To make diets containing the PSMs from an individual *A. rubra* tree, we mixed 10 g of the base diet with 3.3 mL of the prepared PSM extract. In total, we generated one control diet and 20 PSM-imbued diets, 10 of which represented the local trees along the Pysht River and 10 of which represented the non-local trees along the Little Hoko River. While diets were still warm, we loaded each diet into three randomly selected well locations of each 96-well microplate. Microplates were allowed to cool overnight to ensure diets were solidified before deployment. While loading microplates with diet, we determined the initial mass of diet added to each well by recording the mass of each plate after the addition of diet into each individual well.

***In-situ river mesocosm construction and deployment***

Flow-through mesocosms were constructed from rectangular plastic containers modified with 64 mm mesh lining on all sides, including the lid and the bottom of each container. Six mesocosms were deployed in a row across the channel width of the Pysht River and were anchored in place using stones from the riverbed. We used stones that had been brushed of visible periphyton in order to prevent larvae from having an alternative food source to the supplied diet plates. Further, none of the diets included in our *in-situ* river mesocosms contained antibiotics, as these were only added to disrupt the gut microbiome prior to the feeding assay.

To ensure that similar sized larvae were added to each mesocosm, we measured initial length and width of the larval case prior to transferring larvae to their assigned mesocosm. We monitored mesocosms every 2-3 days to estimate the amount of diet remaining. Care was taken to remove plates before any individual well was completely consumed to prevent underestimates in diet consumption. To record final mass, diet pellets were removed from individual wells using a metal pick, gently dried with a paper towel, and weighed.

***Statistical analyses for feeding assays***

To evaluate the effects of the antibiotic treatment and occurrence of *A. rubra* PSMs on the quantity of each diet consumed, we used a linear mixed effects model with a composite variable as a fixed effect. This composite variable was generated by combining both the gut microbiome treatment, which included the categories of control versus antibiotic-treated larvae, and the diet treatment, which included the categories of the control diet, diets imbued with local *A. rubra* PSMs and diets imbued with non-local *A. rubra* PSMs. Therefore, this composite variable included six unique categories for use in planned orthogonal contrasts, including: 1) antibiotic-treated larvae fed local diets, 2) control larvae fed local diets, 3) antibiotic-treated larvae fed non-local diets, 4) control larvae fed non-local diets, 5) antibiotic-treated larvae fed control diets and 6) control larvae fed control diets. To control for variation attributed to each individual tree used to prepare the *A. rubra* diets, we included tree identity as the random effect term. The linear mixed effects model was built using the *lmer* function of the lmerTest package.[12] We completed an arcsine square root transformation of diet mass consumed to meet assumptions of normality. Using this model, we performed planned-orthogonal contrasts by calculating the estimated marginal means for each contrast with the *emmeans* and *contrast* function of the emmeans package.[13] First, we tested the following planned contrast: the effects of the gut microbiome on consumption of the control diet vs. all *A. rubra* diets. Second, we tested the following set of planned orthogonal contrasts: the effects of the gut microbiome on consumption of: (1) local diets prepared with the PSMs of trees from the Pysht population, (2) non-local diets prepared with the PSMs of trees from the Little Hoko population, and (3) control diets containing no *A. rubra* PSMs.

***Dissection of* Dicosmoecus *larvae and morphological characteristics***

We measured case width (cm), case mass (g), larval abdomen length (cm), larval total body length (cm), and larval body mass (g). For the case width measurement, the case stabilizer stones located toward the opening of the case were used as a reference point to maintain consistency among measurements. During handling, we kept larvae over ice and blotted them dry to minimize water weight prior to recording case and body mass. For abdomen length, the measurement was taken from the hind legs of the metathorax to the posterior end of the abdomen after the anal claw.

To further minimize contamination, we carried out all dissections in a ThermoScientific 1300 Series A2 biological safety cabinet using UV-sterilized equipment. In brief, larvae were placed into 70% ethanol for 1 minute, transferred to 15% bleach for 1 minute, and then submerged in 0.2 μm nylon filtered-sterilized milliQ water for 2 minutes to remove any residual ethanol or bleach from the larval external surface. As the gut was mostly liquified due to freezer preservation, we penetrated the larval abdomen with a sterile 10 μL pipette tip to remove gut material. We mixed the gut sample with 800 μL of 0.2 μm nylon filter-sterilized PBS and then homogenized by vortexing for 2 minutes. We then passed the gut homogenate through a 13 mm 0.22 μm PES membrane filter, immediately snap froze filters in liquid nitrogen, and stored at -80°C.

***Characterization of the larval gut microbiome via 16S rRNA sequencing***

DNA was extracted from filters using the Qiagen DNeasy PowerSoil Pro Kit with the following modifications. For cell lysis and homogenization, we used a BioSpec bead beater set to 3400 rpm for 45 seconds for two rounds. To prevent DNA degradation, we placed PowerBead Pro tubes over ice for one minute after each round. Additionally, we added 2 x 25 μL of Solution C6 to the center of the filter membrane with two rounds of 5-minute elutions prior to the final elution centrifugation step. To describe the bacterial component of larval gut microbiomes, we targeted the hypervariable V4 region of the 16S rRNA gene using the 515f and 806r primers: 5’-GTGYCAGCMGCCGCGGTAA-3’ and 5’-GGACTACNVGGGTWTCTAAT-3’, respectively [14,15]. Libraries were prepared and sequenced at Argonne National Laboratory. Samples were multiplexed with a 12 bp tag and then sequenced on an Illumina MiSeq instrument set to a PE150 read cycle.

Raw DNA sequences were processed with QIIME2 v.2022.8. We used the DADA2 plugin to quality filter, trim primers, denoise reads (--p-pooling method = “pseudo”), remove chimeras, and merge paired end reads, generating Amplicon Sequence Variants (ASVs). ASVs were taxonomically classified using the SILVA 138 16S rRNA database.[16,17] We then used ASVs to build a phylogenetic tree by aligning sequences with MAFFT v.7.505 [18] and infer relatedness among ASVs through a generalized time-reversible CAT model of rate heterogeneity with FastTree v.2.1.11.[19] We then combined the ASV table, assigned taxonomic data, phylogenetic tree, and metadata into a single R object using the phyloseq v.1.36.0 package in R v.4.4.1.[20] To remove any potential host-derived organelle sequences, we removed all ASVs that were assigned as “Eukaryota” and “Unassigned” at the Domain level; “Unassigned” at the Phylum level; “Chloroplasts” at the Order level; and “Mitochondria” at the Family level using the *subset_taxa* function. We also filtered out singletons and ASVs associated with three PCR blanks that were sequenced on the same MiSeq run using the *prune_taxa* function.

***Statistical analyses for amplicon data***

To test for differences in antibiotic treated and control larval gut microbiomes, we utilized distance-based redundancy analyses (db-RDA) where antibiotic treatment was a main effect. We generated the Bray-Curtis distance matrix using the *vegdist* function within the vegan package v.2.6-8.[21] UniFrac distance matrices were constructed using the *UniFrac* function within phyloseq. Each model was built using the *dbrda* function within vegan. To infer significant differences between treatment groups, we then used a permutational analysis of variance for each of our db-RDAs (nperm = 10,000). To determine how antibiotics affected alpha diversity of *Dicosmoecus* gut microbiomes, we used a Hill numbers approach using the *d* function within vegetarian v.1.2.[22,23] We chose to analyze alpha diversity at orders *q* = 0, 1, and 2, which represent ASV richness, the exponential of Shannon’s Diversity and the inverse of Simpson’s Diversity, respectively. For each value of *q,* we created three separate linear models where alpha diversity was predicted by antibiotic treatment as a main effect. For each model an analysis of variance was used to determine significant differences. This approach allowed us to compare the effects of our antibiotic treatment on abundant and rare bacterial taxa of the gut microbiome. We evaluated beta diversity with betadisper, a method that is similar to a Levene’s test in multivariate space, which analyzes the homogeneity of group dispersion of community composition.[24] We built a linear model where a pairwise distance matrix using distances from each observation to the group centroid was predicted by treatment. An analysis of variance was used to determine significant differences in beta diversity between antibiotic-treated and control larval gut microbiomes.

In addition, to broadly characterize bacterial composition of the gut microbiome between antibiotic-treated and control *Dicosmoecus* larvae, we generated a heat map for the top 10% most abundant bacterial families with the *comp_heatmap* function (sample seriation = “Identity”) of the microViz package. We then compared taxonomic differences between treated and control larval gut microbiomes via edgeR.[25] We first obtained proportional abundances and then applied an abundance threshold of 0.0001 to filter out low abundance ASVs using the *transform_sample_counts* and *prune_taxa* functions. Proportional abundances were then normalized with the *calcNormFactors* function (method= “RLE”). To indicate differential abundances between our control and antibiotic-treated groups we used the *exactTest* function, which calculated log-fold change values per ASV. Significant differences in ASV abundance were determined after a Benjamini-Hochberg false discovery rate correction (α = 0.001).

The potential functional consequences of treating gut microbiomes with antibiotics was assessed with a standard PICRUSt2 (v. 2.5.3) pipeline.[26] We then inferred gene family abundances by running hidden-state predictions for Kyoto Encyclopedia of Genes and Genomes (KEGG) orthology (KO) per genome with the R package *castor* v.1.8.2.[27] Metagenome predictions were generated by normalizing ASVs by predicted 16S copy numbers and their corresponding functional predictions per sample. We then implemented a differential abundance analysis to determine significant differences in predicted KEGG pathways. This was similarly conducted as described above, however with the following modifications. Per sample KO abundances were converted to KEGG pathway abundances with the *ko2kegg_abundance* function in the R packages *ggpicrust2* v.1.7.3 [28] with an abundance threshold set to 0.00001. After a Benjamini-Hochberg FDR correction, 165 KEGG pathways were significant (α = 0.05). These pathways were annotated with the *pathway_annotation* function in *ggpicrust2*, where KEGG pathway identifiers were queried against the KEGG database generating pathway classes, descriptions, and maps. Functional inferences were drawn from closely related taxa, as indicated by an average Nearest Sequenced Taxon Index (NSTI) of 0.22 ± 0.014 SE among the 2886 ASVs retained after discarding 9 ASVs with NSTI values above the 2.0 cutoff. Finally, we subset this data for “Metabolism” pathways at the class level and examined the predicted gene content of the top ten most significant metabolism pathways.

**SUPPLEMENTAL TABLES**

**Table S1.** **Environmental factors of the sites along the Pysht and Little Hoko Rivers where *A. rubra* trees and leaf metabolomes were sourced for the diet assay of this study.** For further context, the Pysht River serves as the local site as the *Dicosmoecus* larvae were collected, treated, and deployed back in mesocosms at ELK 1. Average temperature, pH, conductivity and dissolved oxygen were measured with a Hach Meter HQ2100 from the center of the water column at the center of each site’s river width.

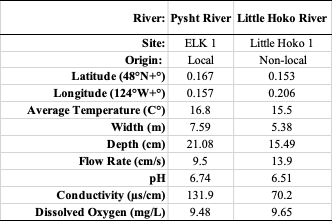

**Table S2. The composition of PSMs within the *A. rubra* leaf metabolome.** Plant secondary metabolites (PSMs) were recovered via tandem liquid chromatography-mass spectrometry from fresh leaf tissue of *A. rubra* trees of the Little Hoko and Pysht Rivers. PSMs were annotated by SIRIUS [29,30] and are listed in the first column followed by the number of features and the median feature intensity for each of these superclasses. We sorted the below table by the median feature intensity to highlight the most prominent PSM superclasses found within the *A. rubra* metabolome. For example, phenylethanoids (C6 – C2), diarylheptanoids, and histidine alkaloids are on average the top three most intense PSM superclasses. Further, the PSM superclasses with the greatest number of unique features are Phenolic Acids (C6 – C1) that contain ellagitannins (n = 315), triterpenoids (n = 202), amino acid glycosides (n = 123) and flavonoids (n = 121). Feature intensities were normalization by the internal standard, sulfachlorpyridzaine.

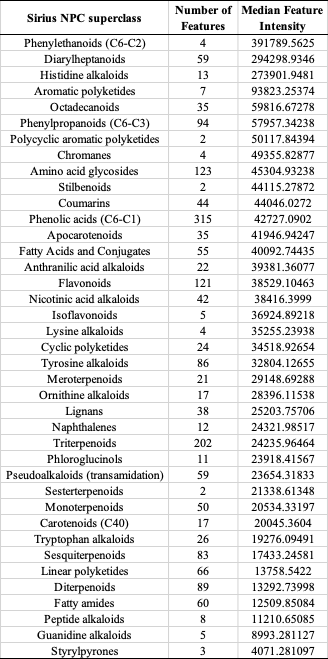

**Table S3. Variable Importance in Projection (VIPs) for Partial Least Squared Discriminant Analyses (PLS-DA) of *A. rubra* leaf PSMs.** Here, we display the top ten VIP features from each model, followed by the mass of each parent molecule in positive mode (m/z), retention time (RT), VIP scores for PLS-DA Component’s 1 and 2, the group each VIP contributes to the most, and the SIRIUS superclass annotation and corresponding zodiac score. Ellagitannins were nested within the phenolic acids (C6 – C1) SIRIUS superclass and were manually verified by parent molecular mass and MS2 diagnostic fragmentation patterns.

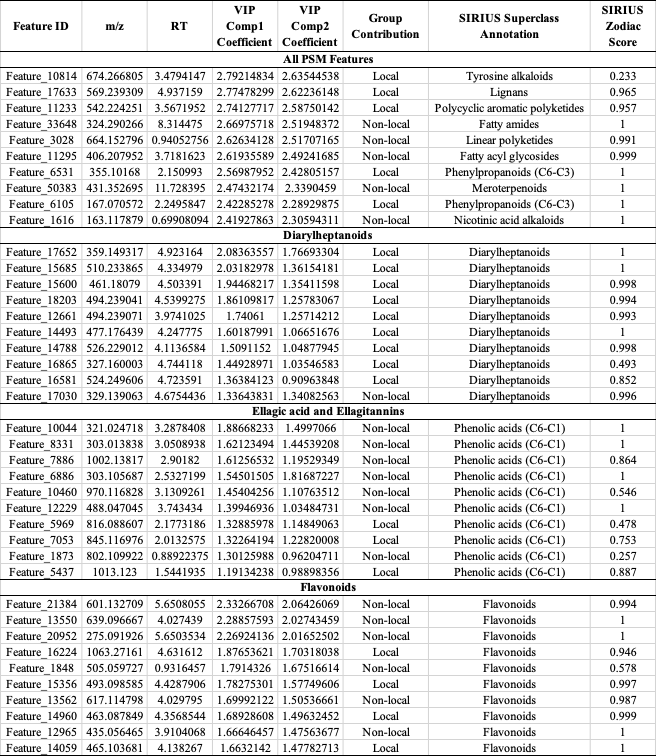

**SUPPLEMENTAL FIGURES**

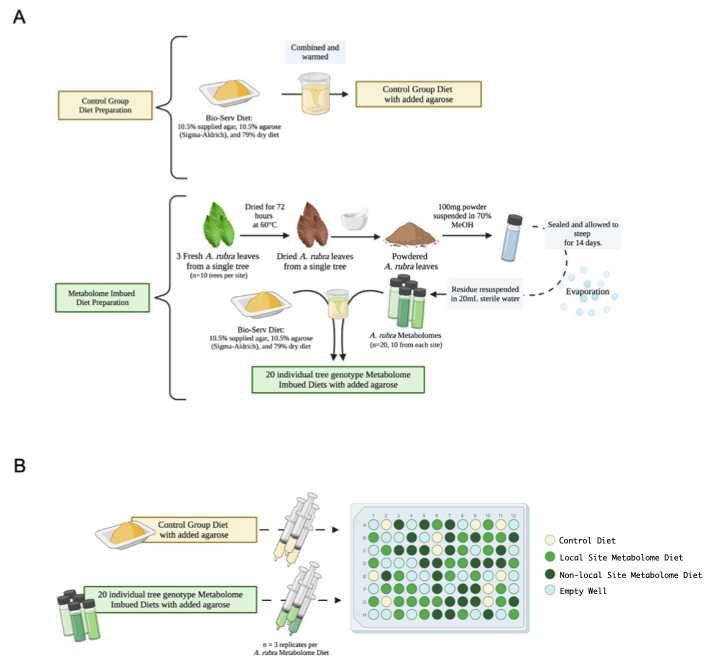

**Figure S1. Methodological approach to prepare control and *A. rubra* PSMs-imbued diet plates. (A)** Control diets were prepared using Bio-Serv diet, provided agar, and additional agarose. PSM-imbued diets were prepared by extracting metabolites from dried *A. rubra* leaves from each of the 10 trees at local and non-local sites following a field-adapted 70% methanol extraction protocol that mirrors our extraction protocol for tandem liquid chromatography-mass spectrometry. **(B)** These diets were then randomly assorted in six 96-well microplates and allowed to solidify before deployment in flow mesocosms at the local site of the Pysht River.

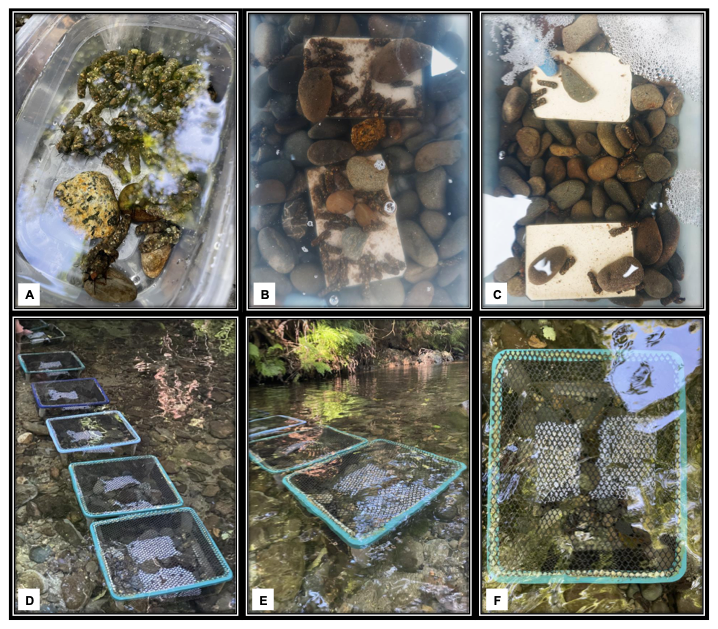

**Figure S2. Images of *Dicosmoecus* larvae during field collection, through antibiotic treatment, and images of the *in-situ* river mesocosms as the feeding assay began. (A)** Collection of *Dicosmoecus* larvae from the Pysht River. **(B – C)** At the Olympic Natural Resources Center, larvae were added to mesocosms with aquarium air stones, river stones from the Pysht River, upside-down 96-well microplates filled with diet, and water from the Pysht River. Larvae randomly assigned to the antibiotic treatment group received diets containing antibiotics and water from the Pysht River that was routinely replenished with antibiotics. **(D)** All six *in-situ* river mesocosms assembled at the ELK 1 site in the Pysht River. **(E – F)** After assembly, *in-situ* flow-through mesocosms remained closed and were monitored every 2-3 days until approximately half of the diets were consumed across all mesocosms.

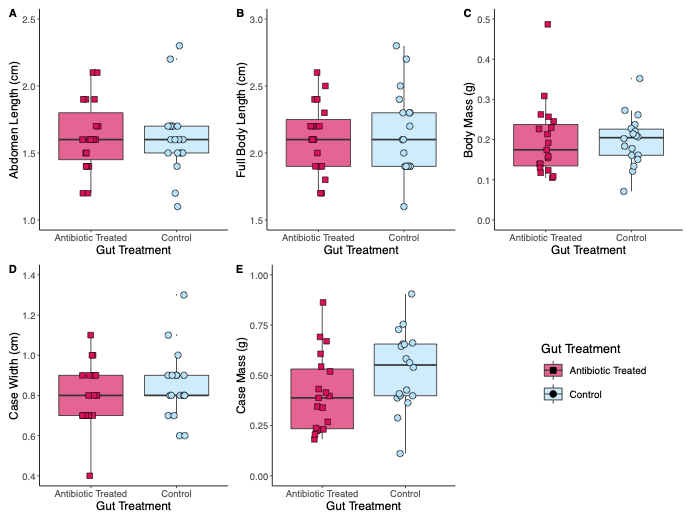

**Figure S3. Individuals of *Dicosmoecus* larvae assigned to the antibiotic-treated versus control groups did not differ in their morphology. (A)** Abdomen length (*F*_1, 35_ = 0.01, *p* = 0.92), **(B)** full body length (*F*_1, 35_ = 0.13, *p* = 0.72), **(C)** body mass (*F*_1, 35_ = 0.045, *p* = 0.84), **(D)** case width, measured as the distance between the two case stone stabilizers near the opening to the case (*F*_1, 35_ = 1.06, *p* = 0.31), **(E)** and case mass (*F*_1, 35_ = 3.35, *p* = 0.076). This suggests that both treatment groups were comprised of individuals with similar distributions in morphological traits and age classes.

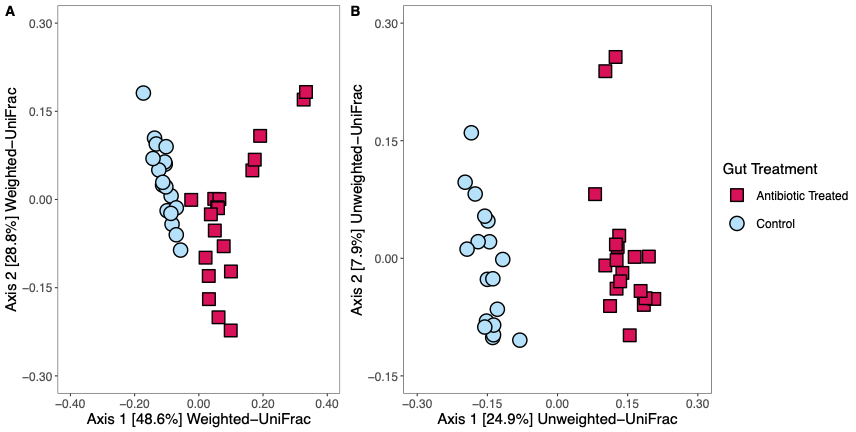

**Figure S4. The gut microbiomes of antibiotic treated and control larvae differ in their phylogenetic membership.** (**A)** abundance-weighted UniFrac (db-RDA— *F*_1, 35_ = 21.91, *p* < 0.001) and **(B)** abundance-unweighted UniFrac (db-RDA— *F*_1, 35_ = 11.06, *p* < 0.001). Each distance-based redundancy analysis (db-RDA) was analyzed with a permutational ANOVA (nperm = 10,000).

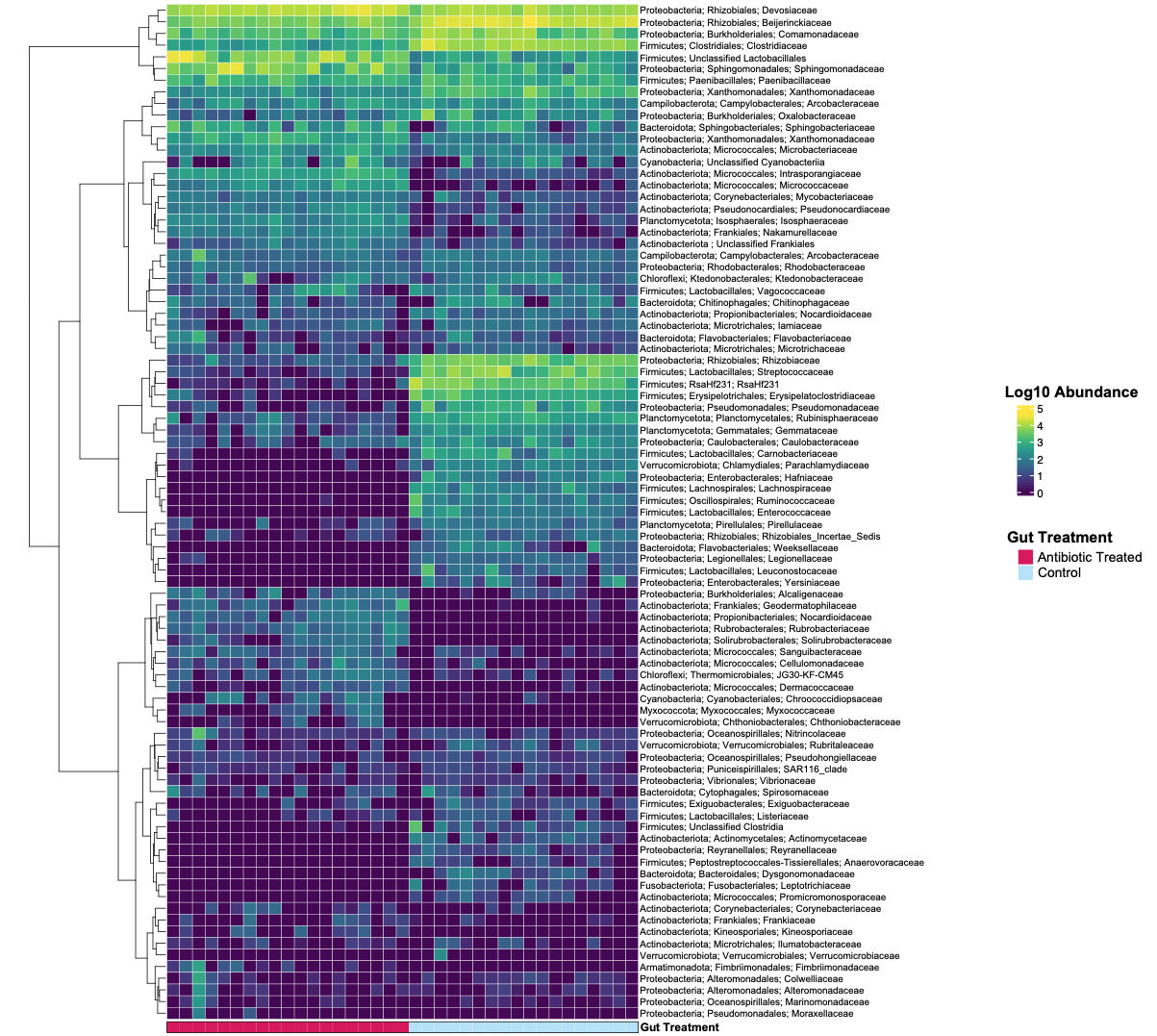

**Figure S5. Caddisfly larval gut microbiomes are enriched in different bacterial taxa between antibiotic treatment groups.** This heat map represents bacterial log 10 abundances at the family level for larval gut microbiomes treated with and without antibiotics (Control). Each column corresponds to a single sample, whereas rows are the bacterial phyla, order, and family. On the left side of the heat map, a dendrogram depicts the relationship between sample occupancy of each bacterial family. We pruned our ASV table by applying an abundance threshold as to display the top 10% most abundant bacterial taxa found within treated and control gut microbiomes (Number of ASVs = 291). Here, we aim to provide a broader characterization of taxonomic shifts in bacterial families found within the gut microbiomes of control and antibiotic treated caddisfly larvae.

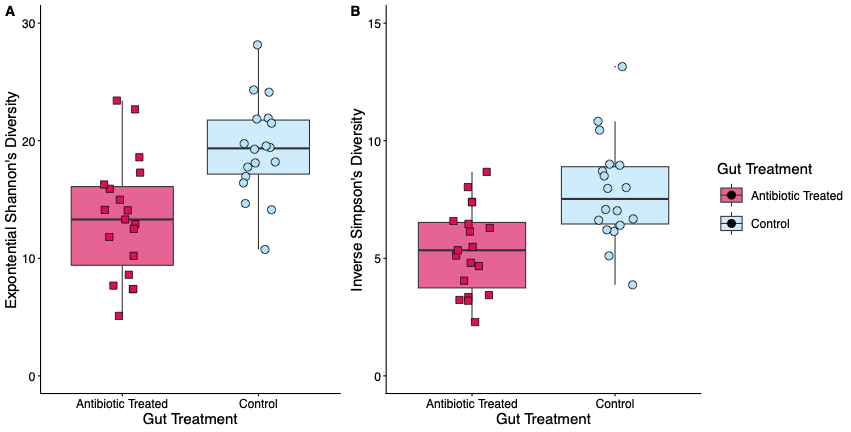

**Figure S6. Bacterial communities of the gut microbiome were more diverse in control larvae in comparison to antibiotic treated larvae. (A - B)** Shannon Diversity and Simpson’s Diversity among control versus antibiotic-treated individuals (Shannon’s Diversity: *F*_1,35_ = 15.11, *p* < 0.001, inverse Simpson’s Diversity: *F*_1,35_ = 13.67, *p* < 0.001).

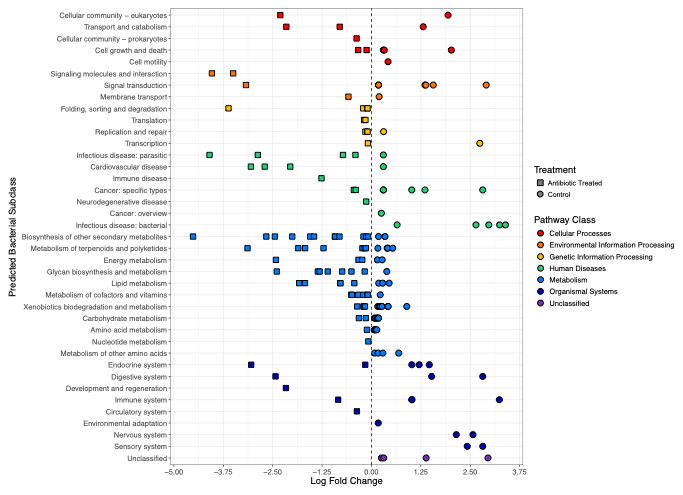

**Figure S7. Predicted bacterial KEGG pathways of the gut microbiome differs between antibiotic-treated and control *Dicosmoecus* larvae.** Antibiotic-treated individuals were differentially under-represented in a broad array of predicted functional pathways ranging in classes from cellular processes, metabolism, and organismal systems. Antibiotic-treated *Dicosmoecus* larvae also experienced an over-representation in many metabolic pathways. Positive log-fold change values represent bacterial predicted functional pathways that are over-represented in control *Dicosmoecus* gut microbiomes relative to antibiotic-treated individuals. Negative log-fold change values represent functional pathways that are over-represented in antibiotic-treated gut microbiomes relative to control individuals. Differential abundance analysis was implemented using edgeR (p < 0.05 after false discovery rate correction; Robinson et al., 2009).

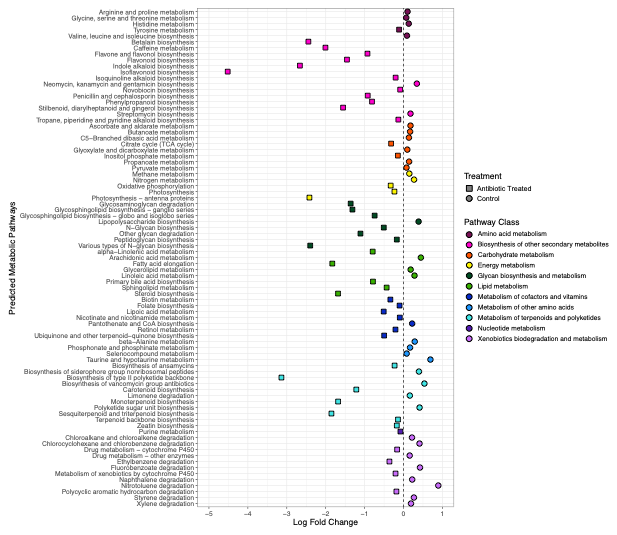

**Figure S8. Honing in specifically on pathways involved in metabolism, predicted bacterial pathways of the gut microbiome are altered after treating *Dicosmoecus* larvae with antibiotics.** KO abundances were converted to KEGG pathway abundances and a differential abundance analysis was implemented using edgeR (p < 0.05 after false discovery rate correction). To determine how antibiotic treatment changed predicted metabolic function within the caddisfly gut microbiome, we illustrate only those pathways under the category “Metabolism.” Positive log-fold change values, plotted as circles, represents bacterial predicted functional pathways that are over-represented in control *Dicosmoecus* gut microbiomes. Negative log fold changes, plotted as squares, represents functional pathways that are over-represented in antibiotic-treated gut microbiomes.

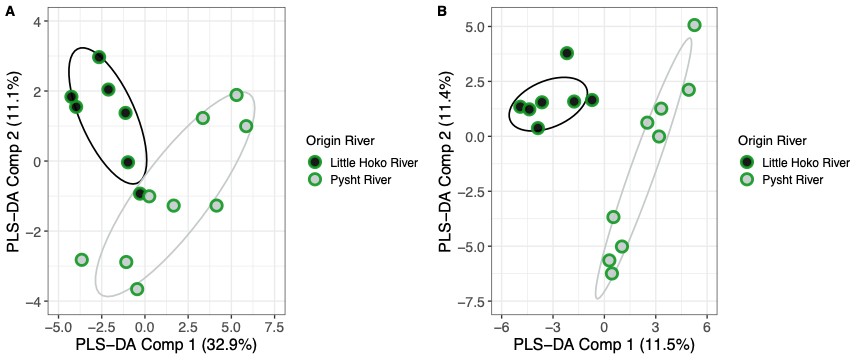

**Figure S9 The ellagitannin and flavonoid fraction of the leaf metabolome vary in their composition among local and non-local populations of *A. rubra*. (A)** Of the ellagitannins (n = 42 features) found within the fresh leaf, we observed distinct clustering via PLS-DA between trees originating from Pysht (local) and the Little Hoko (non-local) Rivers. **(B)** Out of features that were defined by SIRIUS as flavonoids (n = 121 features), we similarly observed supervised clustering via PLS-DA by Origin River. Ellipsoids represent one standard deviation of the multivariate score distribution.

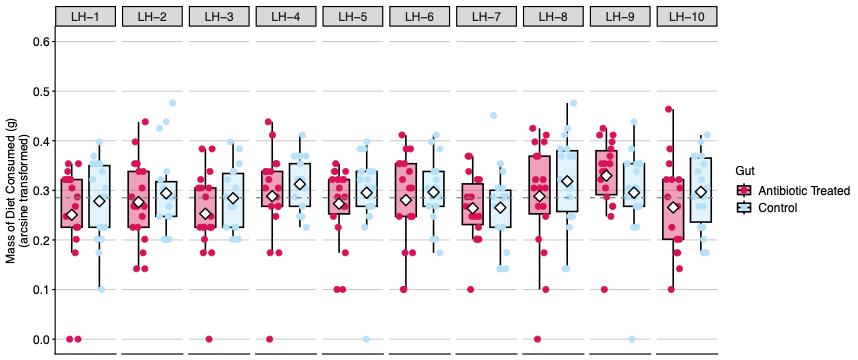

**Figure S10. *Dicosmoecus* larvae with a gut microbiome that has been disrupted by antibiotics display an aversion to diets imbued with PSMs from a non-local population of *A. rubra*.** Compared to control caddisfly larvae that did not receive antibiotics, antibiotic-treated larvae tended to consume less of diets containing the secondary metabolites of non-local *A. rubra* trees growing in the riparian zone of the Little Hoko River. Each of the ten non-local diets were prepared from plant secondary metabolite extracts obtained from individual trees, labelled LH-1 through LH-10. Dashed gray line represents the mean mass of diet consumed across all non-local *A. rubra* imbued diets (displayed in figure as µ = 0.285 ± 0.004 SE; back transformed to milligrams: 85.3mg ± 2.20 SE).

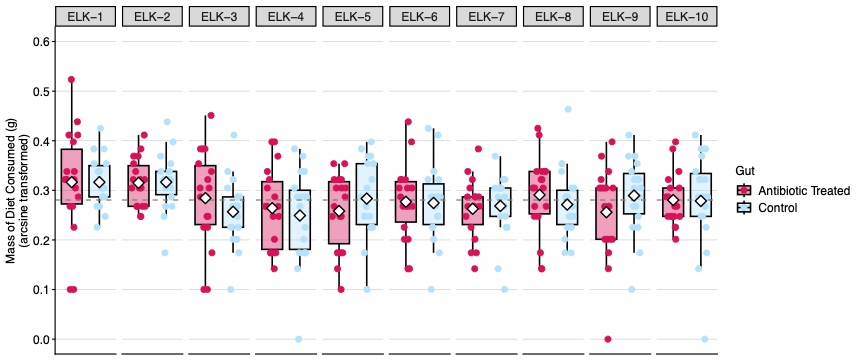

**Figure S11. The state of the *Dicosmoecus* gut microbiome does not significantly alter larval consumption of diets imbued with local *Alnus rubra* PSMs.** Consumption by antibiotic-treated and control caddisfly larvae was similar for ten diets containing the secondary metabolites of local *A. rubra* trees growing in the riparian zone of the ELK study site on the South Fork of the Pysht River. Each of the ten local diets were prepared from plant secondary metabolite extracts obtained from individual trees, labelled ELK-1 through ELK-10. Dashed gray line represents the mean mass of diet consumed across all local *A. rubra* imbued diets (displayed in figure as µ = 0.280 ± 0.004 SE; back transformed to milligrams: 81.3mg ± 2.08 SE).
